# The hypervirulent Group B *Streptococcus* HvgA adhesin promotes brain invasion through transcellular crossing of the choroid plexus

**DOI:** 10.1101/2024.03.26.586743

**Authors:** Eva Aznar, Nathalie Strazielle, Lionel Costa, Claire Poyart, Asmaa Tazi, Jean-François Ghersi-Egea, Julie Guignot

**Author notes:** To whom correspondence should be addressed. Institut Cochin, 22 rue Méchain 75014 Paris, France. CONFLICT OF INTEREST: The authors have declared that no conflict of interest exists.

## Abstract

**Background:** Group B *Streptococcus* (GBS) is the leading cause of neonatal meningitis responsible for a substantial cause of death and disability worldwide. The vast majority of GBS neonatal meningitis cases are due to the CC17 hypervirulent clone. However, the cellular and molecular pathways involved in brain invasion by GBS CC17 isolates remain largely elusive. Here, we studied the specific interaction of the CC17 clone with the choroid plexus, the main component of the blood-cerebrospinal fluid (CSF) barrier.

**Methods:** The interaction of GBS CC17 or non-CC17 strains with choroid plexus cells was studied using an *in vivo* mouse model of meningitis and *in vitro* models of primary and transformed rodent choroid plexus epithelial cells (CPEC and Z310). *In vivo* interaction of GBS with the choroid plexus was assessed by microscopy. Bacterial invasion and cell barrier penetration were examined *in vitro*, as well as chemokines and cytokines in response to infection.

**Results:** GBS CC17 was found associated with the choroid plexus of the lateral, 3^rd^ and 4^th^ ventricles. Infection of choroid plexus epithelial cells revealed an efficient internalization of the bacteria into the cells with GBS CC17 displaying a greater ability to invade these cells than a non-CC17 strain. Internalization of the GBS CC17 strain involved the CC17-specific HvgA adhesin and occurred *via* a clathrin-dependent mechanism leading to transcellular transcytosis across the choroid plexus epithelial monolayer. CPEC infection resulted in the secretion of several chemokines, including CCL2, CCL3, CCL20, CX3CL1, and the matrix metalloproteinase MMP3, as well as immune cell infiltration.

**Conclusion:** Our findings reveal a GBS strain-specific ability to infect the blood-CSF barrier, which appears to be an important site of bacterial entry and an active site of immune cell trafficking in response to infection.

## INTRODUCTION

Group B *Streptococcus* (GBS), also known as *Streptococcus agalactiae*, is a commensal bacterium that colonizes the human gastrointestinal and genital tracts of healthy adults. However, in neonates, GBS is a deadly pathogen being the leading cause of neonatal invasive disease worldwide [1]. Neonatal GBS infections occur predominantly during the first 90 days of life. Early-onset disease (EOD), which appears in the first week of life, results from GBS inhalation during parturition and manifests as respiratory distress, sepsis, and, less commonly, meningitis. In contrast, meningitis is frequent in late-onset disease (LOD). LOD manifests from 7 days to 3 months after birth and is likely to result from mother-to-child post-delivery contamination following ingestion responsible for intestinal colonization [2, 3]. GBS isolates have been classified in capsular types according to their capsular polysaccharide composition (Ia, Ib, II - IX), and in clonal complexes (CC) according to their multi-locus sequence type (MLST). Epidemiological studies have highlighted a remarkable association of capsular type III GBS clonal complex 17 (CC17) with meningitis in neonates. CC17 designated as “hypervirulent” are responsible for 80% to 95% of the cases of neonatal meningitis in both EOD and LOD [2, 4].

GBS meningitis develops following bacteremia [5, 6]. The CNS is separated from the circulation by a system of physiological barriers that blood-borne GBS need to cross to cause meningitis. The microvessels of the brain parenchyma that separate the blood from the brain constitute the blood–brain barrier (BBB) *per se*. The blood-cerebrospinal fluid barrier (BCSFB) constitutes a second physiological barrier which separates the blood from the cerebrospinal fluid (CSF) and is located at the choroid plexus (CP) in the lateral, third and fourth ventricles of the brain. Although CP from different ventricles have different embryonic origins and specific expression patterns, they share a common function in producing cerebrospinal fluid and supporting barrier function [7]. The barrier function of the BCSFB is mainly achieved by choroid plexus epithelial cells (CPEC) that are polarized cells. The presence of tight junctions (TJ’s) and adherent junctions (AJ’s) at the apical level of CPEC contributes to the maintenance of cell polarity and barrier function [8]. BCSFB is also an interface between peripheral and central immune responses and is recognized as an entry site of immune cells into the CNS during both immunosurveillance and neuroinflammation [9]. Like the BBB, the BCSFB may act as a portal of entry for some blood-borne pathogens into the CNS [10]. However, brain penetration *via* the BCFSB is rarely documented. This is due to the few available cellular models of CPEC that have been developed and the technical challenge of infecting CPEC from the basolateral side to follow the pathophysiological process, since *in vivo*, blood-borne pathogens first encounter the basolateral side of CPEC after crossing the choroid plexus capillaries.

Several studies have shown that numerous virulence factors expressed by GBS contribute to the crossing of the BBB [1]. In addition to virulence factors common to most GBS strains, the two CC17-specific surface proteins, HvgA and Srr2, and the highly associated protein, Spb1, play an important role in the unique ability of CC17 strains to cross the BBB by promoting adhesion and internalization into brain endothelial cells. [11–13]. While the host cellular receptor of HvgA remains unknow, Srr2 binds fibrinogen and specifically recognizes the α5β1 and αvβ3 integrins which are overexpressed by brain endothelial cells during the neonatal period [12, 14]. Interestingly, we have shown that these two integrins are also expressed by the cerebral vessels of CP but not of CPEC, and that GBS CC17 can be found in the lateral ventricle outside of the CP capillaries suggesting that a route of infection *via* the BCSFB is likely [12]. This hypothesis is also supported by two other studies *i*) histopathological examination of a fatal case of CC17-associated LOD showed bacteria intimately associated with choroid plexus epithelial cells [11] and *ii*) models of bacterial dissemination in mice have predicted that out of the three BBB interfaces (BBB *per se*, meningeal barrier and CP), the CP is the main interface targeted by GBS for initial transmigration in the brain [15].

In this study, we used *in vivo* experiments as well as inverted cell culture insert system of immortalized and primary CPEC to explore GBS interaction with CP epithelial cells. Our results show that GBS CC17 displays higher capacity to cross CPEC monolayer than GBS non-CC17. Transcytosis of GBS CC17 across CPEC is facilitated by the expression of the HvgA adhesin and occurs by a transcellular mechanism involving clathrin. CPEC cells respond to infection by secreting numerous chemokines. Leukocytes, including T lymphocytes, were observed in the CP of infected mice following infection and may play a role in the neuroinflammatory response to infection. These data reveal that the BCSFB is an important interface during GBS meningitis development, allowing bacterial entry and immune cell recruitment in response to infection.

## METHODS

### Bacterial strains, antibodies and primers

Bacterial strains, antibodies and primers used in this study are listed in Table 1, 2 and 3 respectively. GBS strains were cultured in Todd Hewitt (TH) in standing filled flasks broth or agar (Difco Laboratories, Detroit, MI) at 37°C. Spectinomycin was used at 100 μg.ml^−1^ to maintain GFP expressing plasmid pGU2664 [16].

**TABLE 1:** Bacterial strains used in this study.

| Strains | Characteristics | Reference |
| --- | --- | --- |
| BM110 | CC17 isolate from neonate<br>blood culture (serotype III) | [40] |
| NEM316 | CC23 isolated from neonate<br>blood culture (serotype III) | [41] |
| $\Delta srr2$ | BM110 deleted in *srr2* gene | [42] |
| $\Delta hvgA$ | BM110 deleted in *hvgA* gene | [11] |
| $\Delta spb1$ | BM110 deleted in *spb1* gene | [13] |
| BM110-GFP | BM110 strain carrying pGU2664 plasmid<br>for GFP expression | [12] |

### Animal experiments

A bacteremia-derived meningitis mouse model was used with juvenile (three weeks old) mice. Briefly, mice were challenged by intravenous injection of 2×10^7^ colony forming unit (CFU) of exponentially growing BM110-GFP strain for 4 hours (h) or 24 h of infection, or with 3.5 x10^6^ CFU for 40 h. After infection, mice were deeply anesthetized by intraperitoneal injection of a mix of ketamine/ xylazine (180/20 mg.kg^-1^ body weight) then sacrificed by extensive intracardiac perfusion using sterile phosphate buffer saline (PBS) to remove blood from the circulation followed by 4% paraformaldehyde (PFA) perfusion. Brains were collected and maintained in 4% PFA for 24 h. All animal experiments described in this study were conducted in accordance with the guidelines of Paris Cité University, in compliance with the European animal welfare regulation (http://ec.europa.eu/environment/chemicals/lab_animals/home_en.html) and were approved by the Institut Cochin University Paris Cité animal care and use committee (CEEA34; Agreement number 33235 and 18580).

### Brains slices immunostaining

After fixation in 4% PFA, brains were washed in PBS and then either embedded in agarose for sectioning with a vibratome or immersed in 30% sucrose overnight for cryostat sectioning. Brains embedded in 5% agarose were cut into 500 μm-thick sections using a vibratome (Leica VT1220, Biosystems). Brain sections were clarified using Rapid Clear® CS mounting solution (SunJin Lab) according to the manufacturer’s instructions and then immunostained with anti-CD31 (a marker for capillaries) and anti-transthyretin (TTR, a phenotypic hallmark marker for CPEC). Brains in 30% sucrose were cut into 40 μm-thick sections using a cryomicrotome (HM 450, Thermo Scientific). For long-term storage, slices were kept at −20°C in a cryoprotectant solution (30% ethylene glycol, 30% glycerol in PBS). Sections were analyzed after immunostaining using anti-CD31, anti-TTR and anti-CD45 or anti-CD3 antibody to label total leucocyte or T lymphocyte, respectively. The tissue sections were imaged using an Olympus IXplore Spin confocal microscope controlled by the Metamorph 7.7.5 software. Images were processed using ImageJ.

### Z310 and primary rat CPEC culture conditions

Transformed Z310 is an established cell line derived from rat CPEC and has been described previously [17, 18]. Briefly, cells were seeded on 3 μm pores inverted cell culture inserts (1.12 cm^2^) (Transwell® Corning) precoated with fibronectin-collagen IV (10 μg.ml^-1^) and grown in DMEM containing 10% Fetal Calf Serum (FCS) and 1 μM dexamethasone (Sigma) for 14 days to allow proper differentiation. Cells were routinely tested for mycoplasma contamination (Mycoalert mycoplasma detection kit, LONZA). Primary CPEC were prepared from 2-day-old neonatal rats by the BIP facility (https://www.crnl.fr/en/plateforme/bip) as described previously [19, 20]. Briefly, cells were seeded on 3 μm pores inverted cell culture inserts (0.33 cm^2^) and maintained in culture medium consisting of Ham’s F-12 and DMEM (1:1) supplemented with 10% FCS, 2 mM glutamine and 50 μg.ml^-1^ gentamicin. Prior to infection assays, CPECs were rinsed and incubated in serum- and gentamicin-free medium for 24 h.

### Invasion assay in Z310 cells

Bacterial invasion assays were performed in cellular culture media without FCS at a multiplicity of infection (MOI) of 10 bacteria per cell unless otherwise specified. Briefly, bacteria were grown to the mid-log phase, washed twice with sterile PBS then added to the upper Transwell filter compartment to infect cells by the basolateral side. After 1 h of incubation at 37°C under a 5% CO_2_ atmosphere, monolayers were washed 3 times with PBS, and then treated for 1 h with cellular culture media containing penicillin/streptomycin (Gibco) and gentamicin (200 μg.ml^-1^) to kill extracellular bacteria. The monolayers were washed twice with PBS, and lysed by osmotic choc with sterile H_2_O. Appropriate dilutions were plated on TH agar and CFUs counted. The percent of invasion was calculated as follows: (CFU on plate count / CFU in inoculum) × 100. Assays were performed in duplicate and were repeated at least three times. When specified, results were expressed as normalized to the control condition. To decipher host signaling pathways during GBS CC17 internalization, cells were pretreated with cytochalasin D (2 μM), chlorpromazine (28 μM), dynasore (80 μM), PP2 (20 μM), nocodazole (1 μM), wortmannin (100 nM), MK2206 (2 μM), methyl-β-cyclodextrin (2.5 mM), Y27632 (10 μM) or staurosporine (25 nM) for 1 h prior infection with GBS CC17. Inhibitors were left during the time of infection at the same concentration except for dynasore for which concentration was dropped to 10 μM to avoid bactericidal effect. In these conditions, treatments did not affect bacterial or cell viability (data not shown).

### Dextran permeability assay

Paracellular integrity of cell monolayers was assessed by measuring permeability to 10 kDa [14C]-dextran (MC2002, Campro Scientific) or [14C]-sucrose as previously described [19]. [14C]-sucrose was used to assess permeability after drug treatment (Dynazore). However, as sucrose can be utilized by bacteria, [14C]-dextran was used to assess permeability during bacterial infection.

### Transmigration assay

For the bacterial transmigration assay, CPEC and Z310 were infected as performed for the invasion assay. Translocated bacteria in the bottom compartment of the Transwell® inserts (apical side) were counted after 2 or 3h. Appropriate dilutions of cell culture media from the lower compartment were plated on TH agar and CFUs counted. For some experiments, cells were pretreated for 1 h with dynazore (80 μM) prior infection, then dynazore was maintained at 10 μM during the time of infection. Dynazore did not significantly altered permeability of the monolayer (not shown).

### Immunofluorescent staining and image analysis of infected CPEC

Z310 cells grown in inverted configuration on Transwell® filters were infected for 1 or 5 h with BM110 strain expressing the GFP (Table 1). Cells were fixed in PBS containing 4% PFA for 10 min or in methanol for 4 min at −20°C. Following fixation, cells were permeabilized by incubation in PBS containing 0.1% Triton-X100 for 4 min at room temperature then blocked for at least 1 h at room temperature in PBS containing 10% serum. Primary antibodies (Table 2) diluted in 10% serum-PBS were incubated for 1 h at room temperature. After 3 washes with PBS, filters were incubated for 1 h at room temperature with conjugated-secondary antibodies (Jackson ImmunoResearch Laboratories) diluted in PBS containing 10% serum. DAPI was used to counter-stain nuclei. After 3 washes in PBS, filters were mounted in DAKO fluorescent mounting media (DAKO).

**TABLE 2:** Antibodies used in this study.

| target | clone | supplier | use |
| --- | --- | --- | --- |
| Anti-ZO1 | 339100 | Thermofisher | IF |
| Anti-ZO1 | 402300 | Thermofisher | WB |
| Anti-Claudin-1 | 13050-1-ap | proteintech | IF, WB |
| Anti-Occludin | 331500 | Thermofisher | IF, WB |
| Anti-Clathrin | 610500 | BD | WB |
| Anti-TTR | AB215202 | Abcam | IF |
| Anti-CD31 | AF3628 | R&D system | IF |
| Anti-CD45 coupled 647 | 103123 | Biolegend | IF |
| Anti-GBS | pa1-7250 | Thermofisher | IF |

IF: immunofluorescence; WB: Western Blot

For some experiment requiring to discriminate between extracellular and intracellular bacteria, cells were infected with GFP expressing bacteria. Extracellular GFP-expressing bacteria were counterstained in unpermeabilized conditions with an anti-GBS antibody followed by a secondary rhodamine isothiocyanate-labeled antibody. Under these conditions, extracellular bacteria were green (GFP-labeled bacteria) and decorated with red (anti-GBS labeling), whereas intracellular bacteria were solely green (GFP-labeled bacteria). Cells were counterstained with ZO-1 labelling and DAPI.

Image acquisitions were realized using confocal laser scanning microscopy (Leica DMI6000) coupled to a spinning disk confocal head (YokogawaCSU-X1M1). Images were processed using the ImageJ software. Confocal images shown correspond to a projection of the z-stack.

### RNA isolation and quantitative real-time PCR (qRT-PCR)

RNA extraction from cultured Z310 cells was performed using the Nucleospin RNA kit (Macherey-Nagel) according to the manufacturer’s instructions. RNA was quantified using the NanoDrop ND-100 (ThermoFisher) and cDNA synthesis was performed from 1 µg of RNA using oligo-dT (ThermoFisher) and SuperScript IV RT (ThermoFisher). RT-qPCR was performed using Power SYBR Green PCR system (ThermoFisher) and specific primers (Table 3) on the LightCycler 480 Real-Time PCR System (Roche). Fold change was calculated using the comparative 2^-ΔΔCt^ method. Results were normalized using 18S or actin as housekeeping genes.

**TABLE 3:**
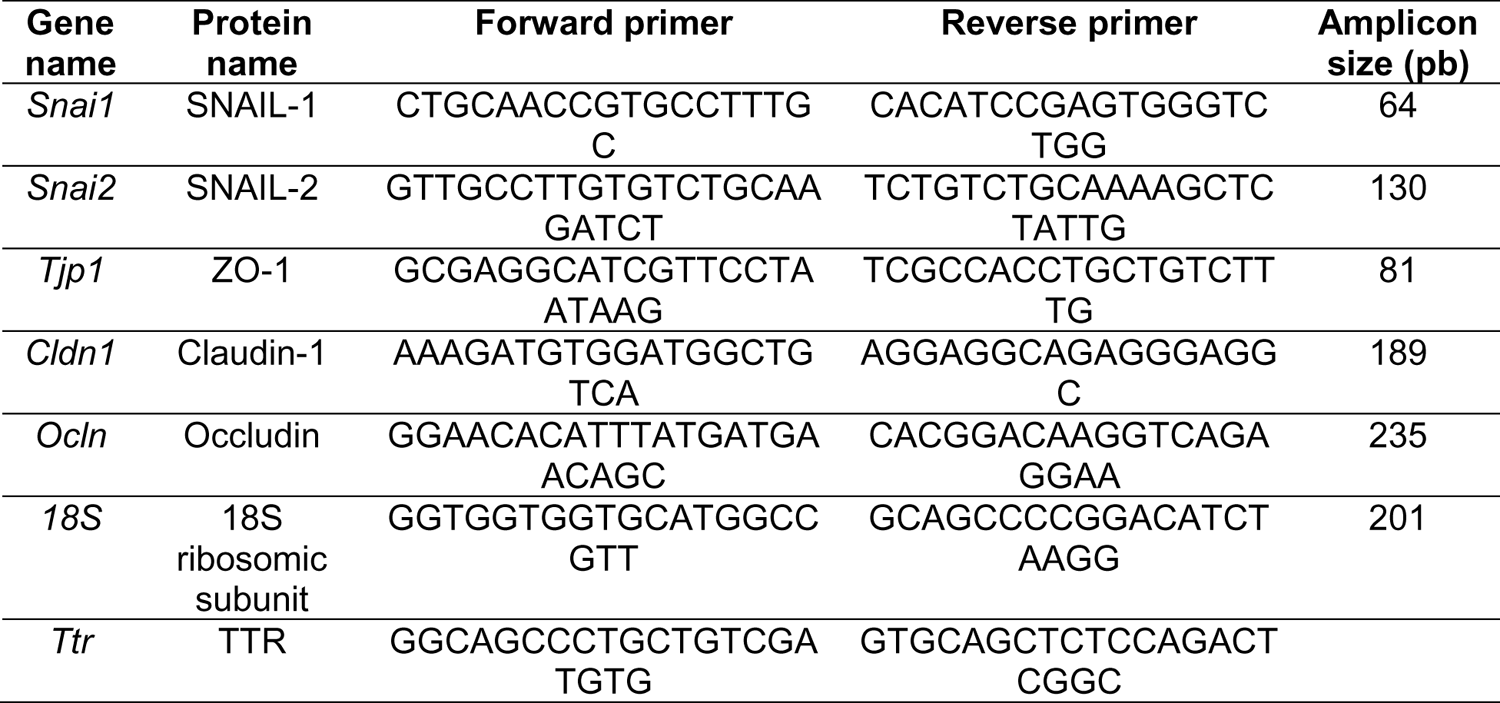
Primers used in this study.

### siRNA transfection

To silence the expression of clathrin, pools of 4 siRNA duplexes (ON-TARGET plus SMARTpool siRNA from Dharmacon) were used. The silencer select negative control-1 siRNA (Ambion) was used as negative control. SiRNA transfection was performed onto undifferentiated Z310 cells. Briefly, cells were trypsinized then transfected with siRNA using the Nucleofactor Kit T (Amaxa Biosystems) following manufacturer’s instructions. Transfected cells were seeded at 20,000 cells/well in 24-well plates for 48 hours, then a second round of transfection was performed with 25 nM siRNA using Lipofectamine RNAiMAX Reagent (Invitrogen) according to the manufacturer’s instructions. Cells were used 24 hours after the second transfection. Efficiency of knockdown was assessed by Western blot analysis.

### Western Blot analysis

For Western Blot, proteins were separated by SDS-PAGE then transferred to PVDF membrane. Membranes were blocked in PBS containing 5% skimmed milk and incubated for 1 h with primary antibodies (Table 2). After washing and incubation with HRP-conjugated secondary antibody (Jackson Immunoresearch West Grove, PA), Western Blots were revealed using the ECL detection system (Perkin-Elmer) and signals were detected using the chemiluminescence imaging system (Fusion, Vilber-Lourmat). Anti-β-actin directly coupled to HRP was used as a loading control for Western Blot.

### Cytokine and chemokine secretion following GBS infection

To identify the cytokines and chemokines secreted in the culture supernatant from the apical or basolateral side of the CPEC monolayer after infection, cells were infected at MOI 10 for 3 h. The media were then changed to media containing antibiotics to kill extracellular bacteria and incubated for a further 21 h. Apical or basolateral culture supernatants from 6 infected Transwell® were collected. Apical and basal culture supernatants from uninfected cells were also collected as controls. All supernatants were stored at −80°C until analysis. Cytokine and chemokines levels were measured using a commercial multiplex immunoassay (Proteome Profiler rat XL Cytokine Array Kit; R&D system) which allows the detection of 79 rat cytokines and chemokines according to the manufacturer’s instructions. To this end, the apical or basal supernatants from the 6 Transwell® inserts were pooled to perform the multiplex array. Membranes from the proteome profiler were developed using imaging system (Fusion, Vilber-Lourmat). Densitometry was performed using ImageJ to determine relative protein levels of each cytokine and chemokine from infected and uninfected conditions.

ELISAs were used to confirm a subset of cytokines identified in the antibody arrays (CCL2, CX3CL1, and Galectin-3 rat ELISA kits, Sigma) using apical or basal supernatants from individual Transwell® inserts from the same experiment than the multiplex array.

### Electron microscopy

Following gentle rinsing with prewarmed PBS, Z310 grown in inverted configuration were fixed in 2.5% glutaraldehyde for 1h, then rinced with PBS and post-fixed for 1h in 1% osmium tetroxide. After dehydrating in increasing concentration of ethanol, filters were immersed in an ethanol:Epon mixture (1:1 vol/vol) for 1h before being transferred to pure Epon. Polymerization was carried out at 60°C for 24h. Sections (90 nm) were prepared, mounted on copper grids and stained with uranyl acetate and lead citrate. Acquisition was performed using a JEOL 1011 transmission electron microscope (JEOL, Japan) with an ORIUS 1000 CCD camera (GATAN, France), operated with Digital Micrograph software (GATAN, France).

### Statistical analysis

All assays were performed at least in three independent experiments with each experiment performed in triplicate. Data represent mean ± SEM and statistical analyses were performed using GraphPad Prism 10 (GraphPad Software, San Diego, California). Significance levels were set at \**p* ≤ 0.05; \*\**p* ≤ 0.01;\*\*\**p* ≤ 0.001.

## RESULTS

### Extravascular GBS are found in the CP of all ventricles

In mice models of GBS meningitis, GBS is able to transmigrate across the blood vessels of the CP and is observed in close contact with CPEC of the lateral ventricle as soon as 1 and 4 h post-infection (p.i.), respectively [12, 15]. The CPs of the different ventricles are of different embryonic origin and, although structurally identical, have specific protein expression profiles [7]. Therefore, we analyzed the ability of GBS to interact with the CP of all ventricles at early and late time points (4 and 24 h p.i.). As neonatal GBS meningitis is strongly associated with CC17 strains, juvenile mice (3 weeks old) were inoculated with a CC17 isolate of GBS (strain BM110, Table1). The whole CP of 5 individual mice were analyzed by confocal microscopy to analyze the presence of bacteria. As expected, extravascular GBS were observed in the CP of the lateral ventricle in all mice (5/5 mice) at 4 h p.i. We also detected extravascular GBS in the CP of the 3^rd^ ventricle (4/5 mice) and the 4^th^ ventricle (3/5mice; Fig. 1, upper panel). At 24 h p.i., GBS were also observed in the choroid plexuses of the lateral (all mice), the 3^rd^ (1/5 mice) and the 4^th^ (3/5 mice) ventricles (Fig. 1, lower panel).

**Figure 1:**
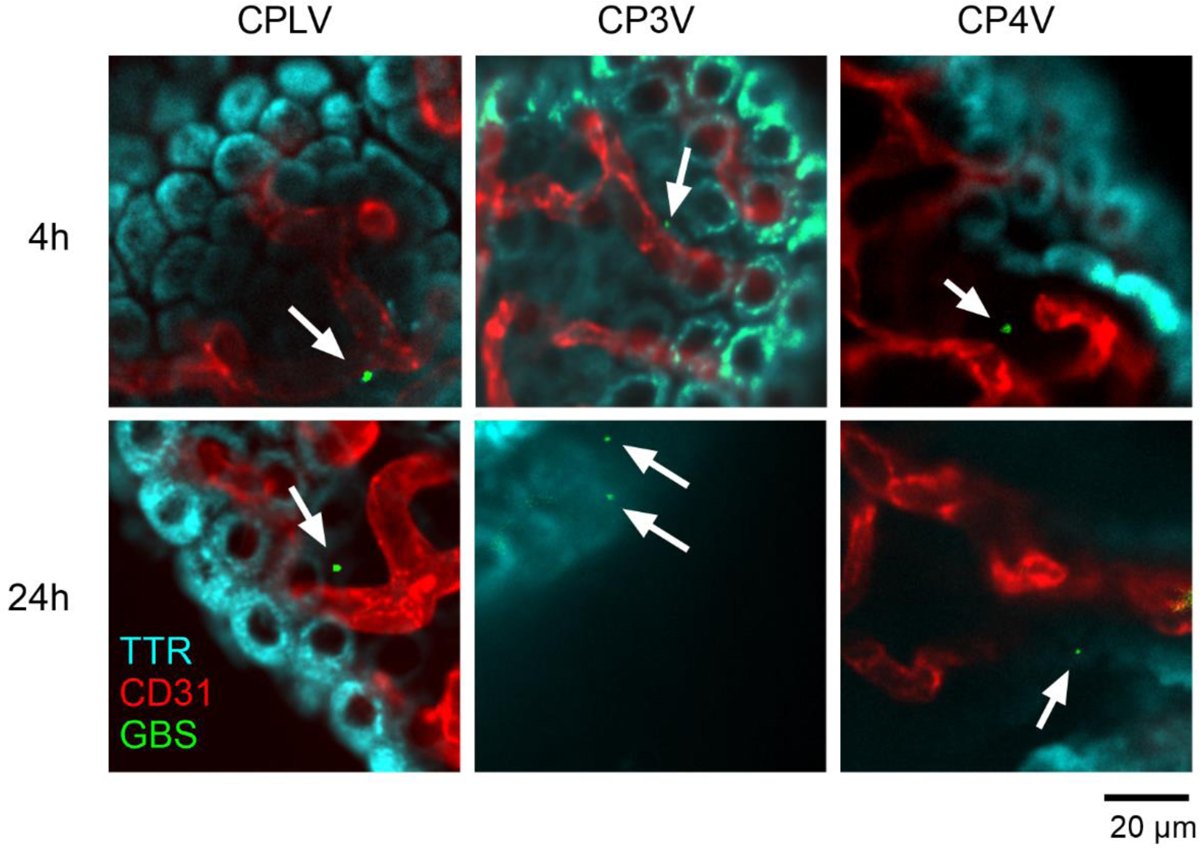
GBS infects CP of all ventricles. Juvenile mice were infected with the BM110-GFP strain (green) for 4 or 24 h. Representative confocal microscopy images of clarified brain sections (500 µm) containing CP from the lateral (CPLV), third (CP3V) and fourth (CP4V) ventricles are shown with CPEC stained with anti-TTR (cyan) and blood vessels with anti-CD31 (red). Scale bar 20 µm.

Taken together, these results indicate that BM110 strain has the ability to migrate across the blood vessels of the CPs of all ventricles as early as 4 h after infection, allowing it to come into contact with the CPECs. GBS were always detected as individual cocci, regardless of the time point. The absence of bacterial aggregates at a later stage (24 h p.i.) suggests that there is no significant bacterial proliferation outside the vessels.

### GBS CC17 infection of CPEC does not disrupt tight junctions

CPECs constitute the main component of the blood-CSF barrier (BCSFB). To characterize their interaction with GBS from a molecular point of view, we used the immortalized Z310 rat cell line described previously [18]. The Z310 cell line expresses the TTR protein, specific transporters (ABC, SLC, MRP) and TJs proteins (ZO-1, Occludin and Claudin-1) and has been described to be a useful model for BSCFB [21]. Since the *in vivo* route of bacterial infection of CP is from the basolateral (blood) to the apical (CSF) side, we set up an inverted Transwell® system to infect fully differentiated Z310 cells from the basolateral side (Fig. 2A). To cross the brain barriers, bacteria must either destabilize cellular junctions to allow their paracellular passage, or be internalized into cells to allow a transcellular passage [6]. In cerebral endothelial cells, GBS induces an upregulation of the transcriptional repressor SNAIL-1 leading to the repression of TJs genes expression and to blood-brain barrier disruption [22]. Therefore, we quantified the expression of the genes encoding SNAIL1, SNAIL2, and TJs proteins (ZO-1, Occluding and Claudin-1) in Z310 cells, after 1 and 5 h of infection by BM110 strain (Fig. 2B and C). Our results show that BM110 infection does not significantly alter the expression of TJs transcriptional regulators or TJs genes. Besides, the protein levels of the TJs proteins ZO1 (220 kDa), Occluding (59 kDa), and Claudin-1 (22 kDa) were also not modified after 1 and 5 h of infection by BM110 (Fig. 2D). Finally, immunofluorescence staining showed that Z310 cells exhibited continuous tight junction strands indicating that TJs proteins are not delocalized after GBS infection (Fig. 2E). Taken together, our results demonstrate that BM110 does not alter the tight junctions of epithelial barrier function during early interactions with Z310 cell line.

**Figure 2:**
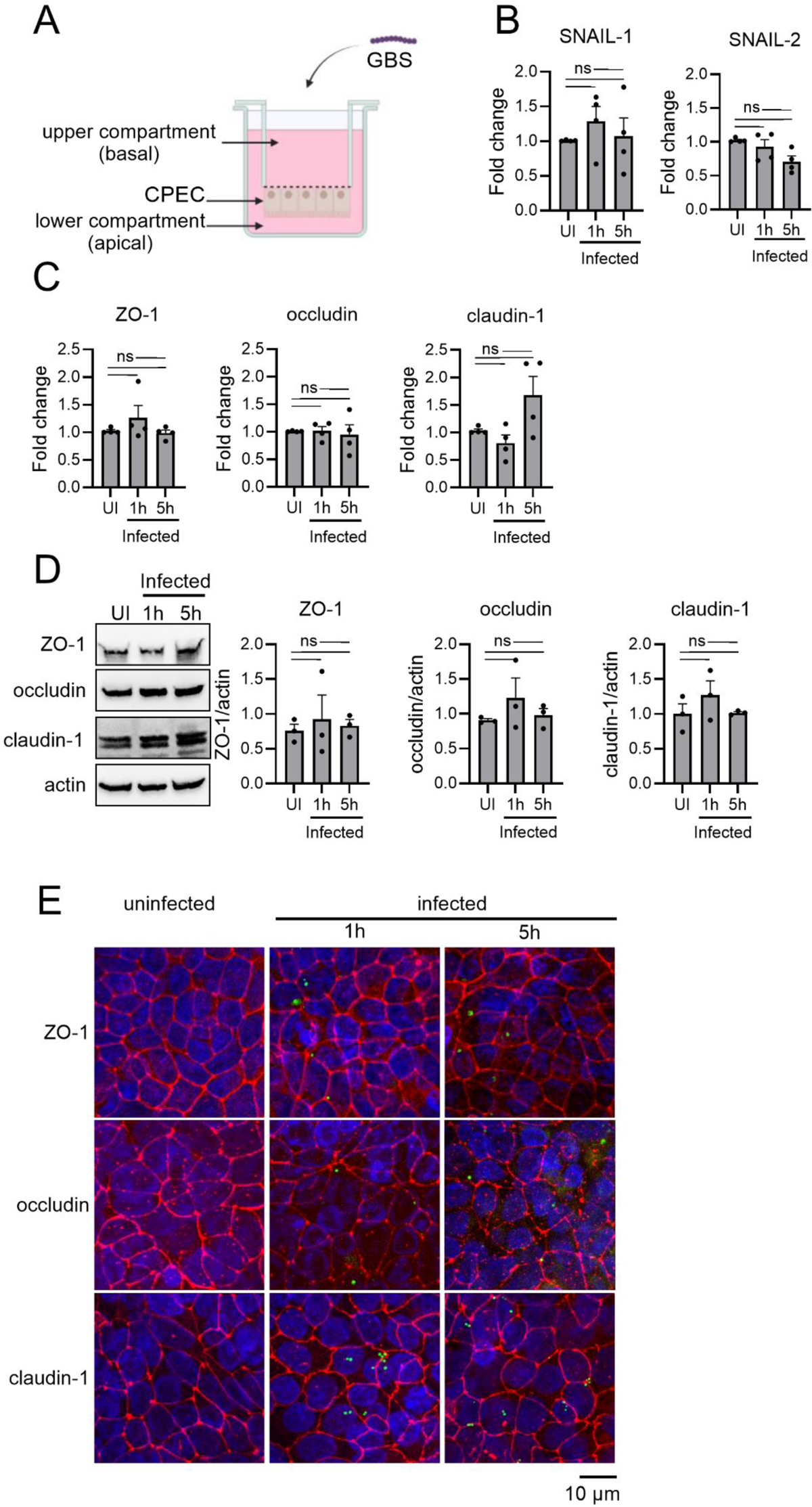
GBS does not affect tight junctions of Z310 choroid plexus epithelial cells. Z310 cells were grown in inverted configuration and infected from the basolateral surface with the GBS strain BM110 at MOI 10 for 1 or 5 h. (A) Schematic representation of inverted culture of Z310 (made by Biorender). (B, C) Fold change in mRNA expression of (B) the tight junction repressors SNAIL-1 and SNAIL-2 and (C) the junctional proteins ZO-1, Occludin and Claudin-1 evaluated by qRT-PCR. (D) Western blot and signal quantification of the ZO-1, Occludin and Claudin-1 proteins normalized over the actin signal intensity. (E) Representative confocal microscopy images of Z310 cells infected with BM110-GFP strain showing the unaltered pericellular distribution of ZO-1, Occludin and claudin-1 proteins. Scale bar 10 µm. Statistical analysis: data shown are mean ± SEM of at least three independent experiments with each dot corresponding to the mean of a triplicate for 1 experiment. (A, B, C) One-way ANOVA with Holm-Sidak’s multiple comparison, with ns, non-significant.

### The HvgA adhesin promotes GBS CC17 internalization in CPEC

We next investigated the ability of BM110 strain to be internalized into Z310 cells after infection. Using differential staining to distinguish extracellular from intracellular bacteria, we were able to detect internalized GFP-bacteria in Z310 cells (Fig. 3A). To gain further insight into the intracellular life of GBS, we performed transmission electron microscopy on Z310 cells (Fig. 3B) and confirmed that the integrity of the cell monolayer and the TJs were not affected after 1 h of infection. In addition, bacteria adhering to the basolateral side of the monolayer as well as bacteria in intracellular vacuoles were observed, suggesting that GBS translocation across the CPEC monolayer may occur by a transcellular mechanism.

**Figure 3:**
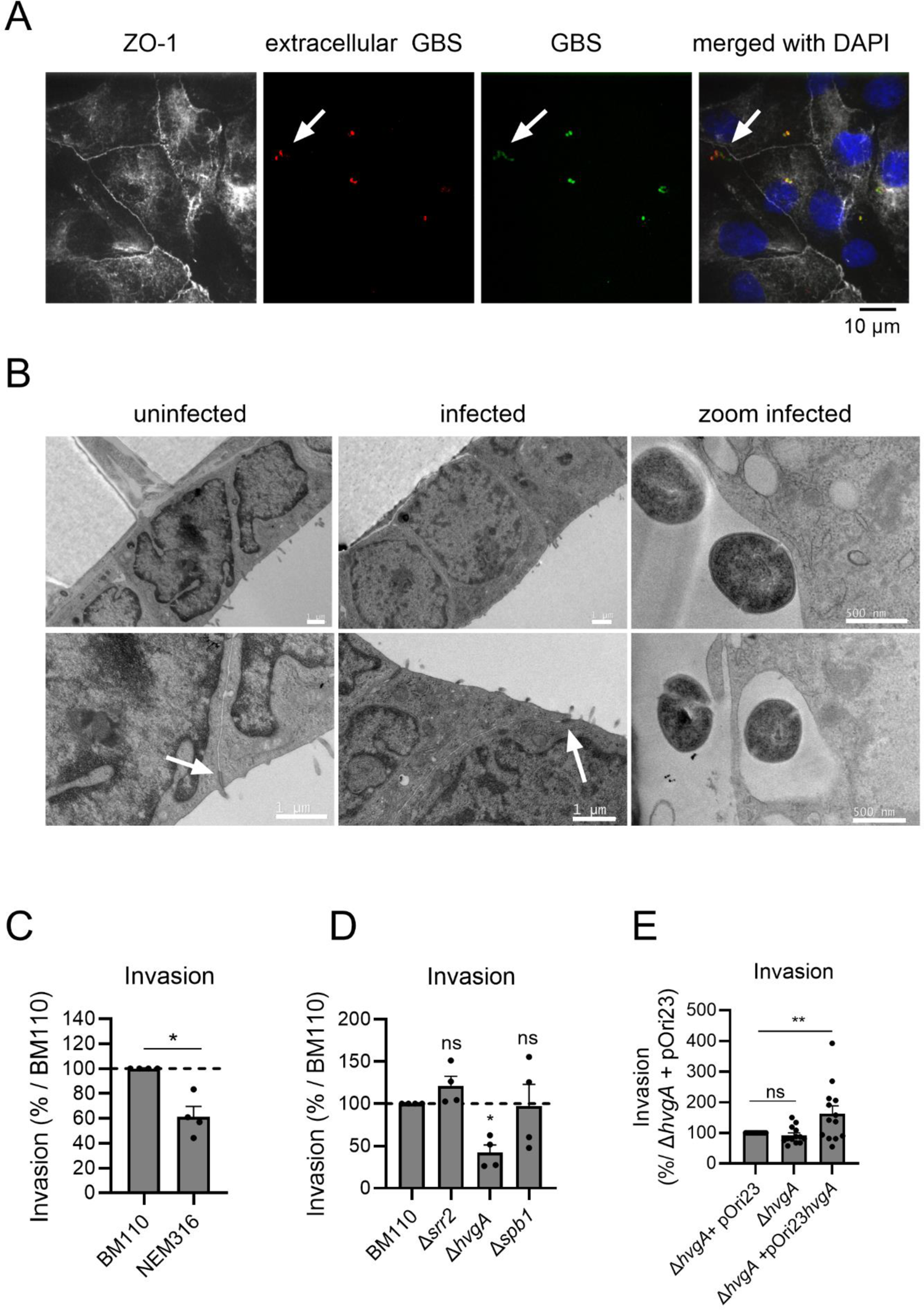
CC17 are internalized in Z310 cells in an HvgA-dependant manner. Z310 cells were grown in inverted configuration and infected from the basolateral surface with GBS strains at (A, C, D, E) MOI 10 or (B) MOI 100 for 1 h. (A) Representative confocal image of Z310 cells realized after a differential staining used to distinguish extracellular GBS (red and green) from intracellular GBS (green only, arrow). Cells were counter-stained with ZO-1 (grey) and with DAPI (blue). Scale bar 10 µm. (B) Transmission electron microscopy micrographs of uninfected (left panel) or infected Z310 (middle panel) showing monolayer and tight junctions’ integrity (white arrows) in control and infected condition (scale bar 1 µm). Bacterial adherence and internalization at the basal surface were observed (right panel, scale bar 500 nm). (C, D, E) Invasion levels of GBS strains were assessed by CFU counts after infection followed by antibiotic treatment to kill extracellular bacteria. Cells were infected with (C) WT BM110 or NEM316 strains, (D) BM110 and its derivative mutant strains (Δ*srr2*, Δ*hvgA* or Δ*spb1*), or (E) with Δ*hvgA* strain, Δ*hvgA* strain carrying an empty vector (+pOri23) or a vector expressing the HvgA protein (+pOri23Ω*hvgA*). (C, D) Results are expressed as the percentage of invasion relative to BM110 strain invasion or (E) to Δ*hvgA* +pOri23 strain. Statistical analysis: data shown are mean ± SEM of at least three independent experiments with each dot corresponding to the mean of a triplicate for 1 experiment. (C) one sample t test, (D, E) One-way Anova with Dunnet’s multiple comparison test with ns, non-significant; *, *p* < 0.05; **, *p* < 0.01.

As CC17 strains have a greater ability to cause meningitis than non-CC17 strains, we next compared the invasion rate of the BM110 strain (GBS CC17) to that of a non-CC17 isolate (NEM316 strain, CC23) (Fig. 3C). CFU enumeration showed that BM110 had a higher ability to invade Z310 cells than NEM316 (mean 1703 CFU versus 1042 CFU, respectively). The neurotropism of GBS CC17 strains has been described to be related to the expression of specific proteins on their surface, namely HvgA, Srr2 and Spb1 [11–13, 23]. We therefore tested the internalization of the corresponding deletion mutants by Z310 cells (Fig. 3D). Our results showed that only the internalization of the *hvgA* mutant was impaired. The introduction into the *hvgA* mutant strain of an expression vector encoding *hvgA* restored bacterial internalization, confirming the involvement of the HvgA adhesin in Z310 invasion (Fig. 3E). In conclusion, our data demonstrate that the specific expression of HvgA by GBS CC17 confers an advantage for CPEC invasion.

### GBS CC17 internalization into Z310 occurs by a clathrin-dependent mechanism and leads to transcytosis

To better characterize the molecular pathways involved in GBS CC17 internalization, we performed invasion assays using a wide array of inhibitors targeting different signaling molecules involved in endocytosis (Fig. 4A). BM110 invasion was inhibited by cytochalasin D, chlorpromazine and dynazore indicating that bacterial internalization by the basolateral side requires actin, the adaptor protein-2 (AP2) and dynamin-2, respectively. In contrast, PP2 nocadazole, wortmannin which inhibit Src kinases, microtubule polymerization and PI3 Kinase, respectively, had no effect. Unexpectedly, MK2206, MβCD, Y27632 and staurosporine which inhibit Akt, lipid rafts, Rock and Protein kinase C considerably increased internalization. The AP2 and dynamin-2 are the major molecular players in clathrin-dependent endocytosis. To confirm that BM110 is internalized by a clathrin-dependent mechanism, we knocked down clathrin expression by siRNA in Z310 cells (Fig. 4B), and observed a significant reduction in BM110 uptake in this condition (Fig. 4C). We next tested whether GBS internalization leads to transmigration across the epithelium (Fig. 4D). To this end, Z310 were infected in the presence or absence of Dynazore, which was very efficient in inhibiting bacterial internalization without affecting the integrity of the TJs. CFU enumeration in the lower compartment showed that BM110 transcytosis was reduced by 81.5% in the presence of Dynazore (mean 153584 CFU versus 28304 CFU, respectively) demonstrating a transcellular passage across Z310 cells. Altogether, our results indicate that GBS internalization into Z310 cells occurs by a clathrin dependent mechanism and leads to bacterial transcytosis.

**Figure 4:**
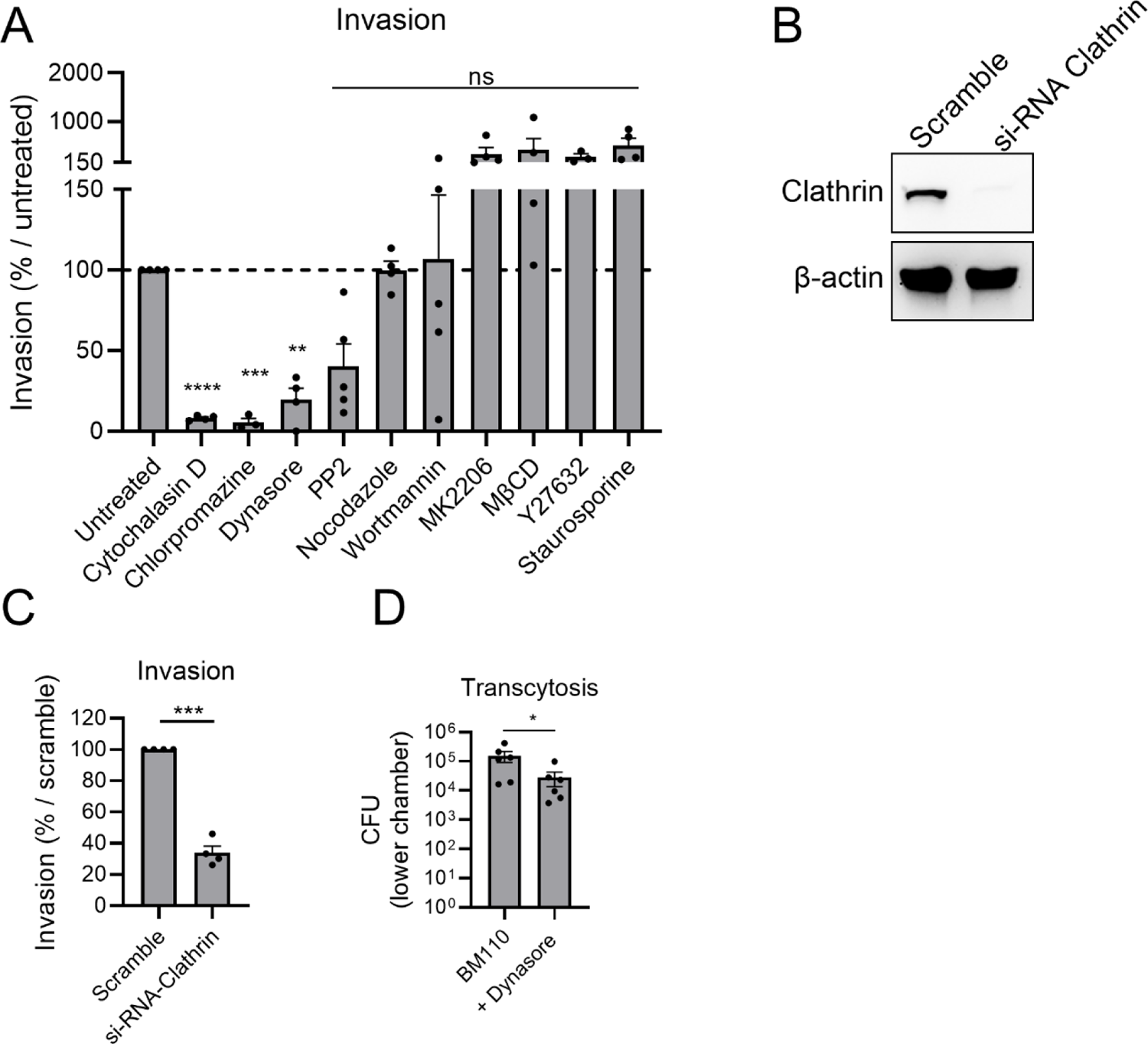
GBS internalization in Z310 cells requires clathrin and leads to transcytosis. Z310 cells were (A, D) grown in inverted configuration to allow infection from the basolateral surface with GBS strains or (B, C) used undifferentiated. (A, C) Invasion levels of BM110 strain were assessed by CFU counts after 1 h infection at MOI 10 followed by antibiotic treatment to kill extracellular bacteria. (A) Cells were pre-treated for one-hour prior infection with cytochalasin D, chlorpromazine, dynazore, PP2, nocodazole, Wortmannin, MK2206, MβCD, Y27632 or staurosporin to inhibit actin polymerization, AP2 protein, dynamin, Src kinase, microtubule polymerization, PI3 kinase, Akt protein, lipid rafts, Rock kinase or protein kinase respectively. Results are expressed as % of bacterial invasion relative to untreated condition. (B) Efficiency of clathrin silencing obtained by siRNA treatment of undifferentiated Z310 cells was evaluated by western blot analysis. (C) Invasion levels of the BM110 strain were assessed by CFU counts after infection of undifferentiated Z310 cells treated by Si-RNA. Results are expressed as % of bacterial invasion relative to scramble siRNA condition. (D) Transcytosis levels of the BM110 strain by Z310 was assessed by CFU counts in the lower compartment after 1 or 2 h of infection following pretreatment or no pretreatment with dynazore for 1h. Statistical analysis: data shown are mean ± SEM of at least three independent experiments with each dot corresponding to (A, C) the mean of a triplicate for 1 experiment (D) the value obtained for one individual filter. (A) One-way Anova with Dunnet’s multiple comparison test; (C) One sample t test (D) Mann-Whitney Test were performed with ns, non-significant; *, p< 0.05 p **, *p* < 0.01; ***, *p* < 0.001.

### GBS CC17 use a transcellular mechanism mediated by HvgA to transcytose across primary neonatal rat CPEC

To consolidate our findings, we next tested the transcytosis of GBS across a primary neonatal rat CPEC monolayer (Fig. 5). Intracellular BM110 was observed in primary CPEC infected from the basolateral side, whereas TJs as assessed by ZO-1 staining were not altered (Fig. 5A). Transcytosis of BM110 and NEM316 across the neonatal primary CPEC was assessed (Fig. 5B). While BM110 transcytosis was detected in 12 out of 20 Transwell® inserts, NEM316 transcytosis was detected in only 8 out of 19. In addition, when transcytosis occurred, the CFU recovered in the lower compartment were higher for the BM110 strain (mean of 18190 CFU) than for the NEM316 strain (mean of 1813 CFU). The paracellular permeability of the monolayer during GBS transcytosis was also tested by measuring radioactive dextran (10 kDa) transfer across the cell monolayers during the transmigration assay (Fig. 5C). The dextran permeability found during BM110 or NEM316 transcytosis was similar to that found in the uninfected condition, indicating that the more efficient transcytosis of BM110 does not involve a differential effect of the bacterial strains on the barrier integrity. We next examined the role of the HvgA protein in BM110 transcytosis (Fig. 5D). CFU recovery in the lower chamber was reduced by 96.6% for the *hvgA* mutant compared to the wild-type strain. Finally, to test whether BM110 transcytosis occurs by a transcellular mechanism, transcytosis was tested in the presence of Dynazore, which strongly inhibits bacterial internalization (Fig. 5E). Under these conditions, no bacteria were detected in the lower compartment (0 out of 9 Transwells) whereas in the absence of dynazore BM110 was recovered in 5 out of 8 Transwells with a mean of 10490 CFU.

**Figure 5:**
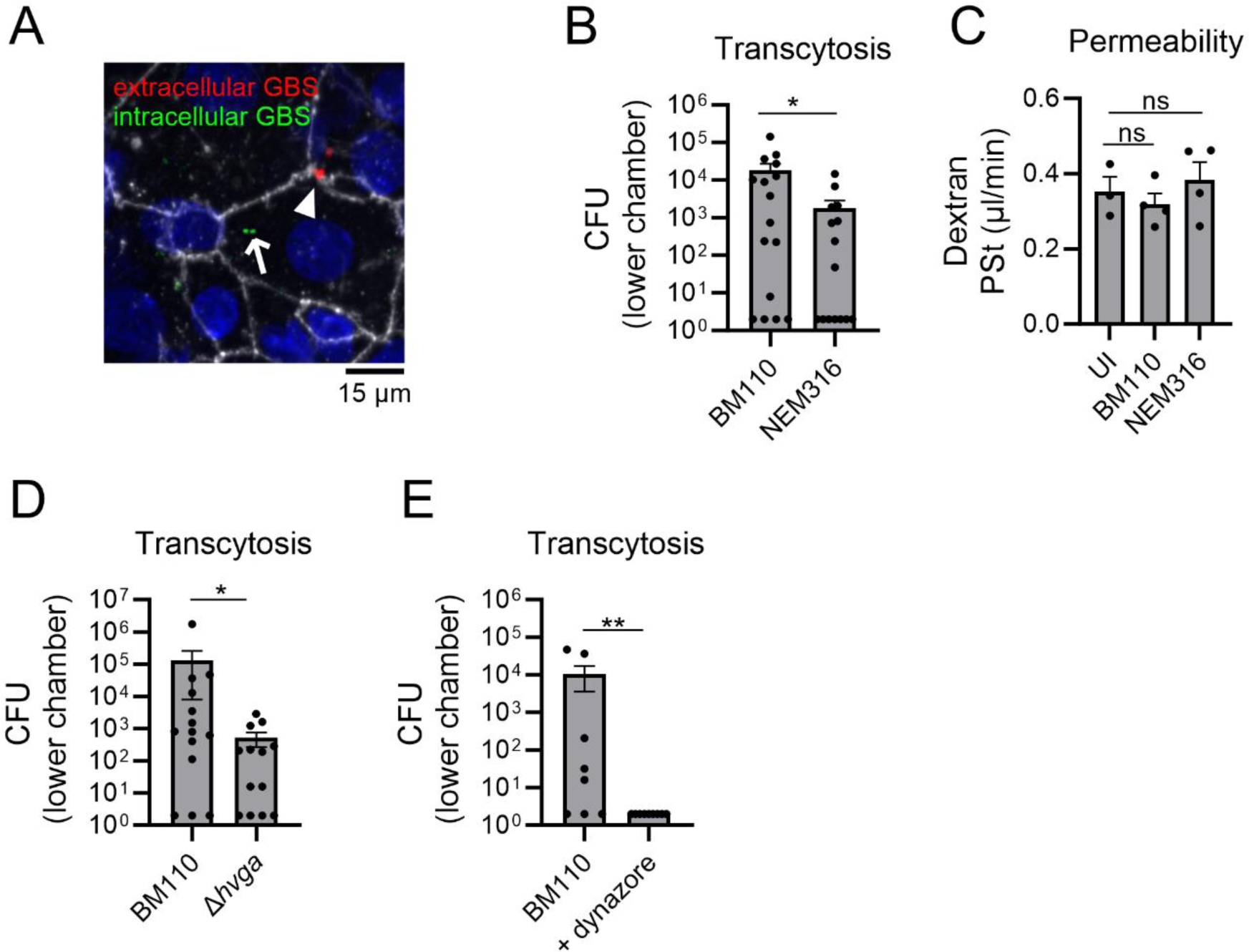
Transcytosis of GBS strains in primary neonatal rat CPEC occurs in an HvgA and clathrin-dependent manner. Primary CPEC were grown in inverted configuration and infected from the basolateral surface with GBS strains at MOI 10 for 2 or 3 h. (A) Representative confocal image of primary CPEC realized after a differential staining used to distinguish extracellular GBS (red and green, arrow head) from intracellular GBS (green only, arrow). CPEC were counter-stained with ZO-1 (grey) and with DAPI (blue). Scale bar 10 µm. (B, D, E) Transcytosis levels of GBS strains across primary CPEC were assessed by CFU counts in the lower compartment after 2 or 3 h of infection (B, D) without pretretament or (E) after pretreatment or not with dynazore for 1h. (C) Permeability of 10 kDa radioactive dextran on primary CPEC was assessed during GBS infection. Statistical analysis: Data shown are mean ± SEM of at least three independent experiments with each dot corresponding to the value obtained for one individual filter. (B, D, E) Mann-Whitney Test and (C) One-way ANOVA with Holm-Sidak’s multiple comparison were performed with ns, non-significant; *, *p* < 0.05; **, *p* < 0.01.

In conclusion, our results show that the BM110 strain, thanks to the expression of the CC17-specific HvgA protein, is more efficient than the non-CC17 strain in crossing the neonatal primary CPEC monolayer that occur by a transcellular mechanism.

### Primary neonatal CPEC respond to GBS CC17 infection by secreting numerous chemokines

We next assessed the cellular response of primary neonatal CPEC that occurs following BM110 infection using a secretome array (Proteome Profiler Rat XL Cytokine Array Kit) to detect specific inflammation-related proteins and their relative expression levels in the supernatant of infected cells. Primary CPEC were infected with BM110 strain for 3 h then treated with antibiotics to kill extracellular bacteria. Apical and basal supernatants from 6 individual Transwell® inserts were collected 24 h p.i., pooled and compared with the uninfected condition. Molecules whose secretion was found to be altered by infection are shown in Fig. 6A and B, while those whose secretion was found unchanged are shown in Fig. S1. Of the 79 rat proteins included in the kit, 13 and 17 were oversecreted in the apical (lower compartment) and basal (upper compartment) supernatants, respectively, after infection (Fig.6 A and B). The chemokines CCL2, CCL3, CCL20, CX3CL1, CXCL2, CXCL5, and G-CSF were the most highly oversecreted proteins in both compartments. In addition, while osteopondin, serpinE1 and galectin-3 were predominantly found in the apical supernatant, MMP3 and cystatin, fibulin, NT3 and IGFBP-2 were predominantly detected in the basal supernatant. These results were confirmed by the quantification of two representative chemokines *e.g.,* CCL2 and CX3CL1, in the supernatants of individual Transwell® inserts using ELISA (Fig. 6C and D). The quantification of CCL2 confirmed a robust and homogeneous over-secretion in the apical and basal supernatants of infected wells compared with control (Fig. 6C). The chemokine CX3CL1 was also homogeneously and highly secreted in the infected apical and basal supernatants, although at lower levels than CCL2 (Fig. 6D). Lastly, we also quantified galectin-3 levels and confirmed that the protein was homogeneously secreted in the apical supernatant, whereas it was barely detected in the basal supernatant (Fig. 6E). In conclusion, GBS CC17 infection elicits a strong immune response by primary neonatal CPEC characterized by the secretion of numerous chemokines.

**Figure 6:**
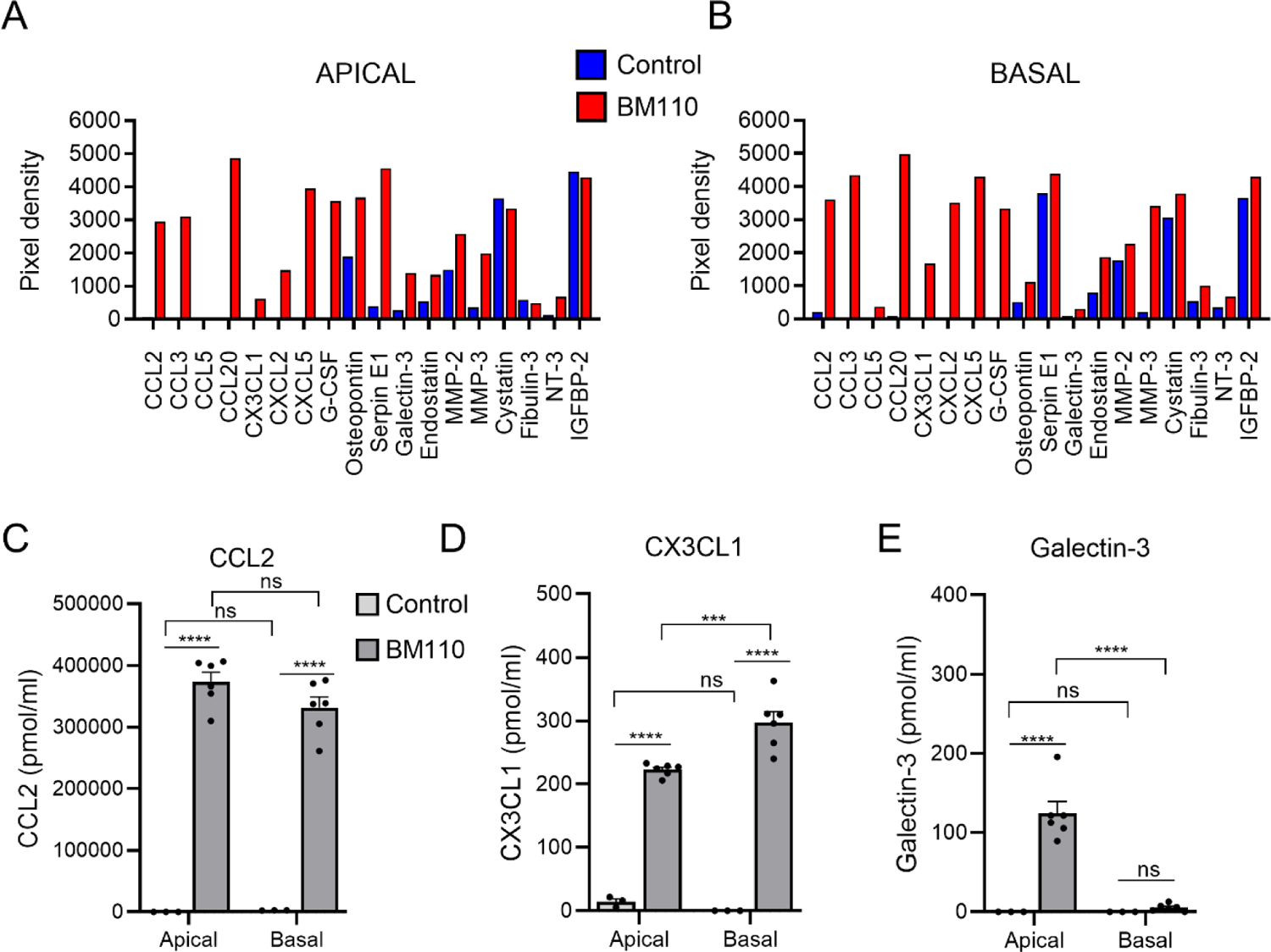
Innate immune response of neonatal primary CPEC induced by GBS infection. Apical and basal culture supernatants from infected primary CPEC were collected and analyzed for the differential expression of secreted molecules following GBS infection. (A, B) pixel intensity quantitative analysis of secreted molecules from the secretome array in the (A) apical supernatant (lower compartment) or (B) basal supernatant (upper compartment). (C, D, E) differential expression of (C) CCL2 (D) CX3CL1 and (E) Galectin-3 were assessed in the apical and basal supernatant of individual wells by ELISA. Statistical analysis: Data shown are mean ± SEM of 6 individual filter with each dot corresponding to the value obtained for one individual filter. (C, D, E) Two-way ANOVA with multiple comparison were performed with ns, non-significant; ***, *p* < 0.001; ****, *p* < 0.0001.

### GBS CC17 meningitis induces leukocytes infiltration in the choroid plexuses

The production of numerous chemokines by primary CPEC following GBS infection prompted us to test whether GBS infection triggered leukocytes infiltration into the CP in our murine model of meningitis (Fig. 7). To this end, mice were infected with GBS BM110 for 40 h and after brain recovery, brain slices encompassing the CP were labelled to observe CPEC (TTR labelling), blood vessels (CD31 labelling) and leukocytes (CD45 labelling) (Fig. 7A). Numerous CD45+ cells adhering to the endothelium and extravascular CD45+ cells were observed in the CP of infected animals compared to uninfected animals. Besides, given that CCL20 is involved in lymphocyte recruitment and is largely secreted by CPECs upon infection, we finally tested if GBS infection was accompanied with T cells infiltration in the CP (Fig. 7B). Immunofluorescence analysis of brain slices revealed that some of the immune cells infiltrating the CP of infected animals were CD3+ cells while no CD3+ cells were observed in the CP of uninfected animals. Altogether, these *in vivo* data and the cytokine array data show that GBS induce a cellular immune response of the CP.

**Figure 7:**
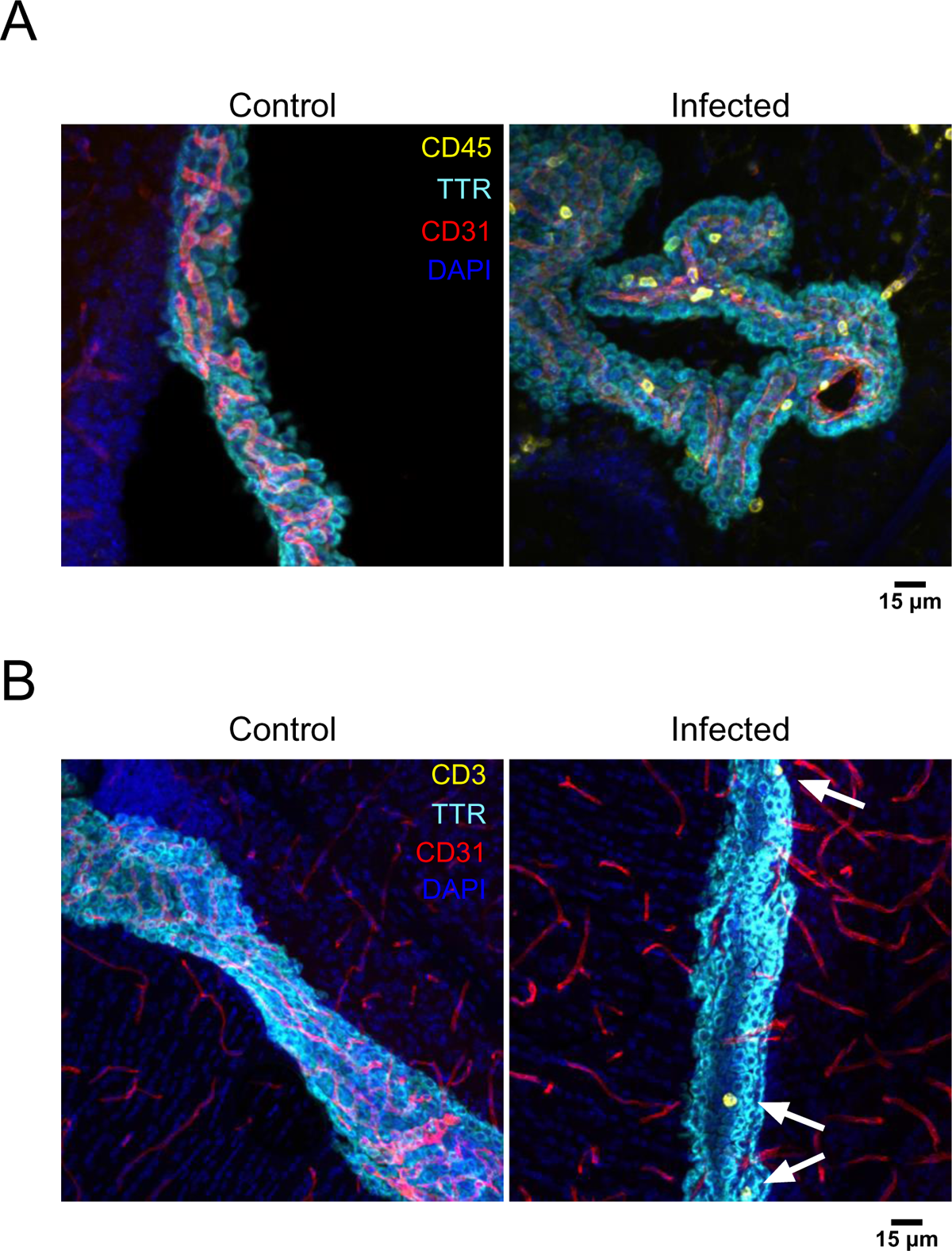
GBS infection promotes leucocytes infiltration in CP. Juvenile mice were infected with the BM110 strain for 40h. Representative confocal image of brain sections (40 µm) immunostained for blood vessels (CD31, red), CPEC (TTR, cyan) and (A) leukocytes (CD45, yellow) or (B) T lymphocytes (CD3, yellow). Scale bar 15 µm.

## DISCUSSION

The hyper-virulent GBS CC17 clone is responsible for the majority of bacterial meningitis in infants under 3 months of age [1]. The aim of this study was to investigate the interaction of GBS with the BCSFB. The results obtained from the observation of mouse brain sections indicate that GBS CC17 has the ability to emerge from the fenestrated blood vessels of the CPs of all the cerebral ventricles. However, whether extravascular bacteria were in the choroidal stroma or within the CPECs, which are the main component of the BCSFB, is difficult to determine due to the limited resolution. In early observations (4 h p.i.), only single cocci and no chains were observed outside the blood vessels. In a mouse model of *Streptococcus pneumoniae* meningitis, only individual cocci, that did not express the cell division protein DivIVA, were able to cross the brain barriers [24]. Our results suggest that this phenotype may be conserved between different *Streptococcus* species. In addition, even at a later stage (24 h p.i.), no bacterial aggregates were detected outside the blood vessels suggesting that there is no significant bacterial multiplication between 4 and 24 h of infection. This is in contrast which what have been observed with other pathogens with meningeal tropism, such as *Neisseria meningitidis*, which develops microcolonies at the cell surface of CPEC [6, 25]. We also observed that the frequency with which extravascular bacteria were observed varied according to the location of the CP in the brain. GBS was systematically observed in the CPs of the lateral ventricles, but its presence in the other ventricles was not systematic. Observations in the CP of the 3^rd^ ventricle could be explained by its surface area, as it is much smaller than the PCs of the lateral ventricles and the 4^th^ ventricle. This does not explain the lower proportion of mice in which bacteria were found in the 4^th^ ventricle, which has a surface area similar to that of the lateral PCs. The difference in the protein expression profile between the PCs of the lateral ventricles and the 4^th^ ventricle could be at the origin of this difference in observation frequency. The lower capacity of GBS to reach the CP of the 4^th^ ventricle may explain the rarity of cerebellar damage associated with GBS infection, whereas rhombencephalitis is more common with others meningeal pathogens such as *Listeria monocytogenes* [26]. Further observations on a larger number of mice, with accurate quantification of extravascular bacteria, would allow statistical analyses to confirm this result. Several meningeal pathogens have been described to alter the junctions of CPEC. Down regulation of TJs or AJs has been observed following *Streptococcus suis* or *Borrelia burgdorferi* infections [27–29]. Using immortalized or primary CPEC, we showed that, in contrast to these pathogens, GBS does not alter CP epithelial junctions. This is also in contrast to what happens at the BBB, where GBS induces TJs proteins repression associated with SNAIL-1 overexpression after 5 h of infection of brain endothelial cells [22]. A recent study suggested that neonatal CPEC junctions are weaker than in adults due to higher Wnt activity [15]. Yet, in this study, expression of TJs genes and their transcriptional regulators was not significantly altered in the CP of GBS infected animals, except for SNAIL-2 and Claudin-5, which were repressed [15]. Since claudin-5 is expressed by endothelial TJs but not by CPEC, this suggested that GBS modifies TJs of the CP endothelium but not those of CP epithelium [15]. Our results confirm that GBS does not repress TJs gene expression of the CP epithelia. Rather, and although not significant, we observed a slight increase in TJs proteins (ZO-1, Occluding and Claudin-1) (Fig. 2B and D), as also observed by Travier *et al*. for Claudin-1 [15]. This slight increase in the expression of junctional proteins could be an adaptive response to infection.

Although the permeability of the CP monolayer was not affected, we observed that GBS was able to invade and transcytose through Z310 and primary CPECs. While this mechanism is conserved between Z310 and primary CPEC the level of transcytosis in primary cells was 10-fold lower and not systematic compared to Z310 cells. This may reflect a lower barrier function property in Z310 than in CPEC. This hypothesis is also supported by experiment in the presence of Dynazore, which was not as effective in blocking bacterial transmigration in Z310 compared to CPEC, suggesting that in Z310 bacteria may translocate through unsealed junctions. Bacterial transmigration in CPEC also revealed that transcytosis is a very limiting step in the pathophysiology of the infection as previously described in brain endothelial cells [30]. Nevertheless, in both models, CC17 strain was more proficient at entering and crossing the CPEC barrier compared to the non-CC17 strain, explaining its successful meningeal tropism. This enhanced ability was associated with the expression of the HvgA adhesin. However, in contrast to the invasion of brain endothelial cells, only the HvgA adhesin was found to contribute to CC17 internalization in CPEC, whereas the Srr2 adhesin did not contribute to this phenotype. This is likely explained by the absence of the Srr2 ligands (fibrinogen, α5β1 and αvβ3) on the surface of CPECs, whereas they are highly expressed by brain endothelial cells [12]. HvgA also promotes adhesion to brain endothelial cells and to phagocytes [11, 31]. We have recently shown that in phagocytes, HvgA adhesin may recognize scavenger receptor(s) to promote bacterial adhesion and subsequent internalization [31]. Since scavenger receptors have a wide distribution and are expressed by phagocytes as well as epithelial and endothelial cells, including in the CNS [32], it is tempting to speculate that HvgA promotes adhesion to host cells by recognizing the same receptors on these different cell types. Further research will be required to confirm this hypothesis.

The internalization of GBS CC17 was shown to be clathrin-dependent and to involve actin, dynamin-2, and the AP2 adaptor. Interestingly, *Neisseria meningitidis* and *L.monocytogenes* internalization into CPEC is also dependent on the dynamin-2 suggesting that clathrin-dependent endocytosis may be a common internalization pathway for pathogenic bacteria during basolateral infection of CPEC [33, 34]. Intriguingly, we found that inhibition of Akt, lipid rafts, Rock and protein kinase C strongly increased bacterial invasion. The reason of this observation remains unclear. One hypothesis is that inhibition of endocytic pathways other than clathrin-dependent endocytosis make available signaling molecules required for GBS internalization.

After internalization within CPEC, we have shown that GBS resides in a cytoplasmic vacuole leading to transcellular translocation across the epithelium. Interestingly, in our secretome array, galectin-3 was found to be secreted exclusively at the apical side. Galectin-3 is a very versatile molecule with complex biological functions including cell adhesion, chemoattraction, inflammation or angiogenesis. Cytosolic galectin-3 is also known to accumulate in structures near internalized bacteria and to be released during vacuole lysis [35]. Its polar secretion at the apical side of CPEC may therefore reflect the delivery of vacuole-bound intracellular GBS into the apical supernatant. Surprisingly, in our secretome array, cytokines were only barely detectable. This may reflect the biphasic pattern of cytokine secretion, characterized by an early phase with rapid release of pro-inflammatory cytokines, followed by downregulation of these inflammatory mediators, as previously described for other pathogens [36]. As in our study, supernatants were collected 24 h p.i., this suggests that we may have missed the cytokine response. In contrast, we were able to detect the production of numerous chemokines, as also observed in response to various pathogens [27, 36, 37]. Among them, CCL3, CXCL2, CXCL5 and G-CSF, were found to be oversecreted in response to GBS infection. These mediators are known to be involved in the recruitment or the differentiation of polymorphonuclear cells (PMN). Pleocytosis is a hallmark of meningitis and PMN are the predominant cell type in GBS meningitis. CCL20, CX3CL1 and CCL2 which are chemoattractants for T lymphocytes, were also found in our screen. Consistent with these findings, we observed an infiltration of T lymphocytes in the CP of infected animals. The BCSFB is recognized as a predominant route of T cell trafficking in the CNS in physiological condition, and can contribute to brain inflammation [20, 38, 39]. Further work to characterize the subpopulation of leukocytes and their activation state as they enter the brain will help to clarify their role in GBS meningitis.

In conclusion, this study provides the first insight into the interaction of GBS with the BCSFB. Our data show that GBS is able to cross this barrier by a transcellular mechanism and highlight the predominant role of HvgA adhesin in brain invasion in CP. Overall, our data may provide relevant insights into the modulation and penetration of blood-CNS barriers by pathogens. This is particularly important as it may lead to the discovery of new therapeutic targets to limit the impact of GBS meningitis.

## AUTHOR CONTRIBUTIONS

JG designed the research and EA, LC, NS and JG performed the experiments. JG, EA, NS and JFGE analyzed the data. AT, CP provided intellectual input and guidance. JG wrote the manuscript and AT, NS and JFGE contributed to finalizing the manuscript. All authors discussed results and reviewed the final version of the manuscript.

## ACKNOWLEDGMENTS

We are extremely grateful to Dr Zheng from Purdue university for the gift of Z310 cells. We thank Matthieu Benard from the animal facility for technical help to infect animals, Alain Schmitt from the PIME core facility for electron microscopy and the Imag’IC core facility of the Cochin Institute.

## Abbreviations

BBB: Blood Brain Barrier

BCSFB: Blood Cerebro-Spinal Fluid Barrier

CC17: clonal complex 17

CP: Choroid plexus

CPEC: choroid plexus epithelial cells

CSF: Cerebro-Spinal Fluid

GBS: group B *Streptococcus*

TJ: Tight junctions

LOD: Late onset disease

EOD: early onset disease.

## SUPPLEMENTARY DATA

**Figure S1:**
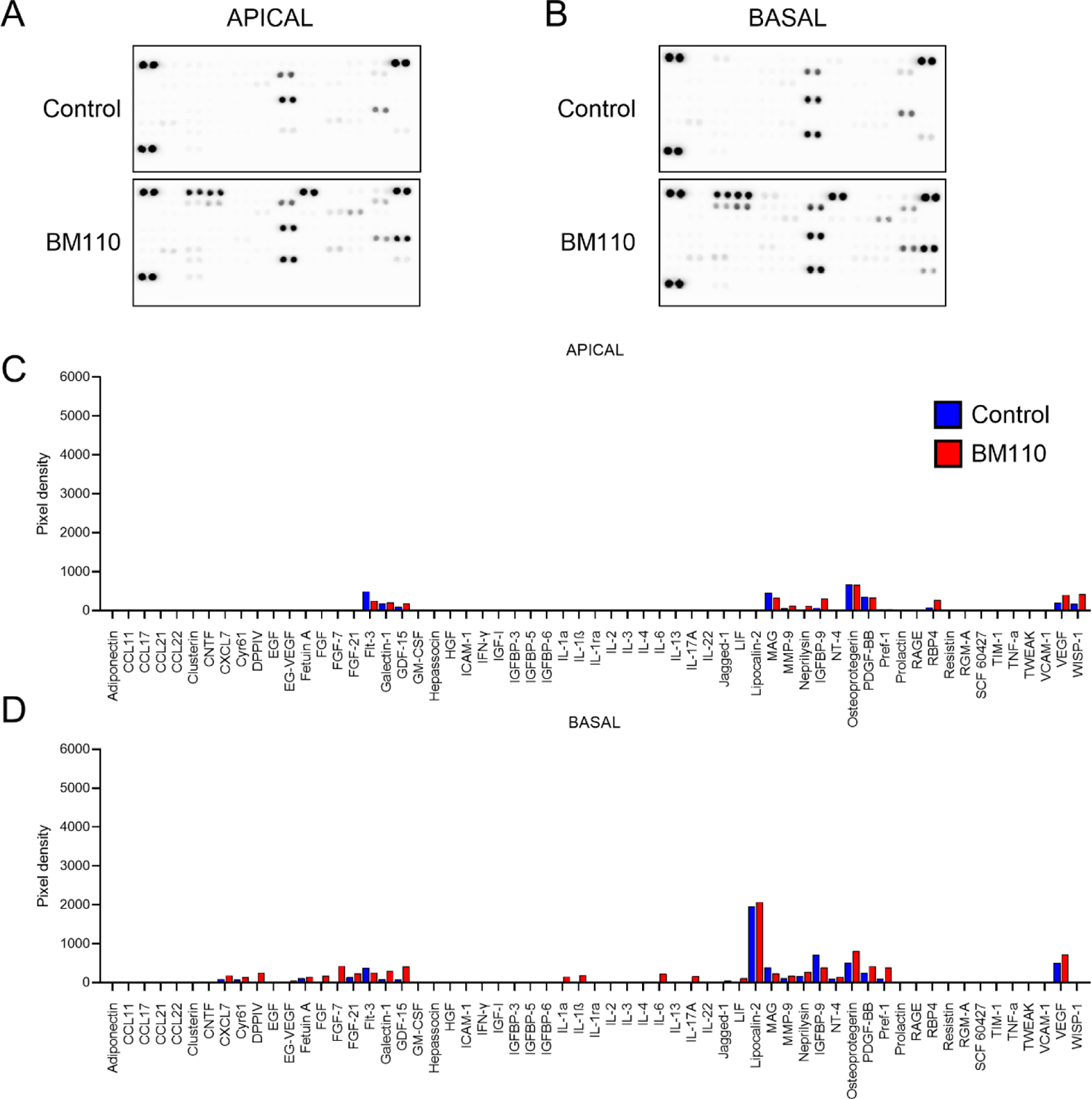
Cytokine array of neonatal primary CPEC following GBS infection. Apical (A, C) and basal (B, D) culture supernatants from infected primary CPEC were collected and analyzed for the differential expression of 79 secreted cytokines associated with GBS infection. (A, B) Results of the membrane cytokines array and (C, D) pixel intensity quantitative analysis for unchanged molecules.

